# Exploring DCAF15 for reprogrammable trageted protein degradation

**DOI:** 10.1101/542506

**Authors:** Seemon Coomar, Dennis G. Gillingham

**Affiliations:** Department of Chemistry, University of Basel, St. Johanns-Ring 19, 4056, Basel, Switzerland

## Abstract

The targeted degradation of proteins by reprogramming E3 ligases with bifunctional small molecules is an exciting area of chemical biology because it promises a set of chemical tools for achieving the same task as siRNA or CRISPR/Cas9. Although there are hundreds of E3 ligases in the human proteome only a few have been shown to be reprogrammable to target new proteins. Recently various arylsulfonamides were shown to induce degradation of the splicing factor RBM39 via the RING type E3 ligase CRL4^DCAF15^. Here we identify the arylsulfonamide most amenable to chemical modifications and demonstrate its behaviour in bifunctional reprogramming.

## Introduction

Over the past few years the use of small molecules to control protein expression levels has emerged as a powerful new therapeutic modality. The importance of controlling cellular proteolysis in transformed cells was illustrated by the approval of proteasome inhibitors such as carfilzomib and bortezomib.^1^ The ubiquitin-proteasome pathway (UPS) is one of the major mechanisms for controlled degradation of proteins. The mechanism^2,3^ involves three ligases (E1, E2, E3) and the key step is the transfer of ubiquitin to a target protein, which is accomplished by the E3 ligase. This last step involves an interaction between the E3 ligase and the protein of interest that is mediated through a substrate receptor (e.g. DCAFs) on the ligase^4^. Recent work has shown that it is possible to reprogram the UPS by binding these substrate receptors with small molecules – a strategy that has led to the development of bifunctional molecules such as proteolysis targeting chimera (PROTACs)^5,6^ as well as molecular glues.^7^ The bifunctional constructs are designed such that one end of the molecule interacts with a novel target protein (neosubstrate) and the other end recruits an E3 ligase for target ubiquitination and degradation. Molecular glues, on the other hand, bind to an E3 ligase substrate receptor and remodel its surface, interrupting some protein-protein interactions, while promoting others.

The list of E3 ligases that seem capable of general reprogramming is short, with only four ligases performing the great majority of all successful PROTACs.^8^ A couple of strategies are being pursued in further expanding PROTACs: using well-established E3 ligase binders and screening and optimizing around the remainder of the small molecule and linker to identify a ternary complex that is active in ubiquitin transfer. A second strategy is to explore new E3 ligases and characterize their reprogrammability^9–13^. With more than 600 putative E3 ligases in the human proteome this strategy is both promising and daunting, since it will require a small molecule binder for every E3 ligase adaptor protein.

The mode of action of a series of antitumor sulfonamides (Fig 1), some of which had even been tested in clinical trials,^14–17^ was only recently clarified in two independent studies.^18,19^ It seems as if these sulfonamides trigger an interaction between the splicing factor RBM39 and the E3 ligase substrate receptor DCAF15 in a molecular glue-type mechanism. This leads to the susequent ubiquitination of RBM39 and its proteolytic degradation. As this mechanism is reminiscent of the more established immunomodulatory drugs (i.e. thalidomides, lenalidomide, pomalidomide), which target CRL4^CRBN^ and have been redesigned as bifunctional degraders, we wondered if the sulfonamides could be repurposed in a similar manner.

**Fig. 1:**
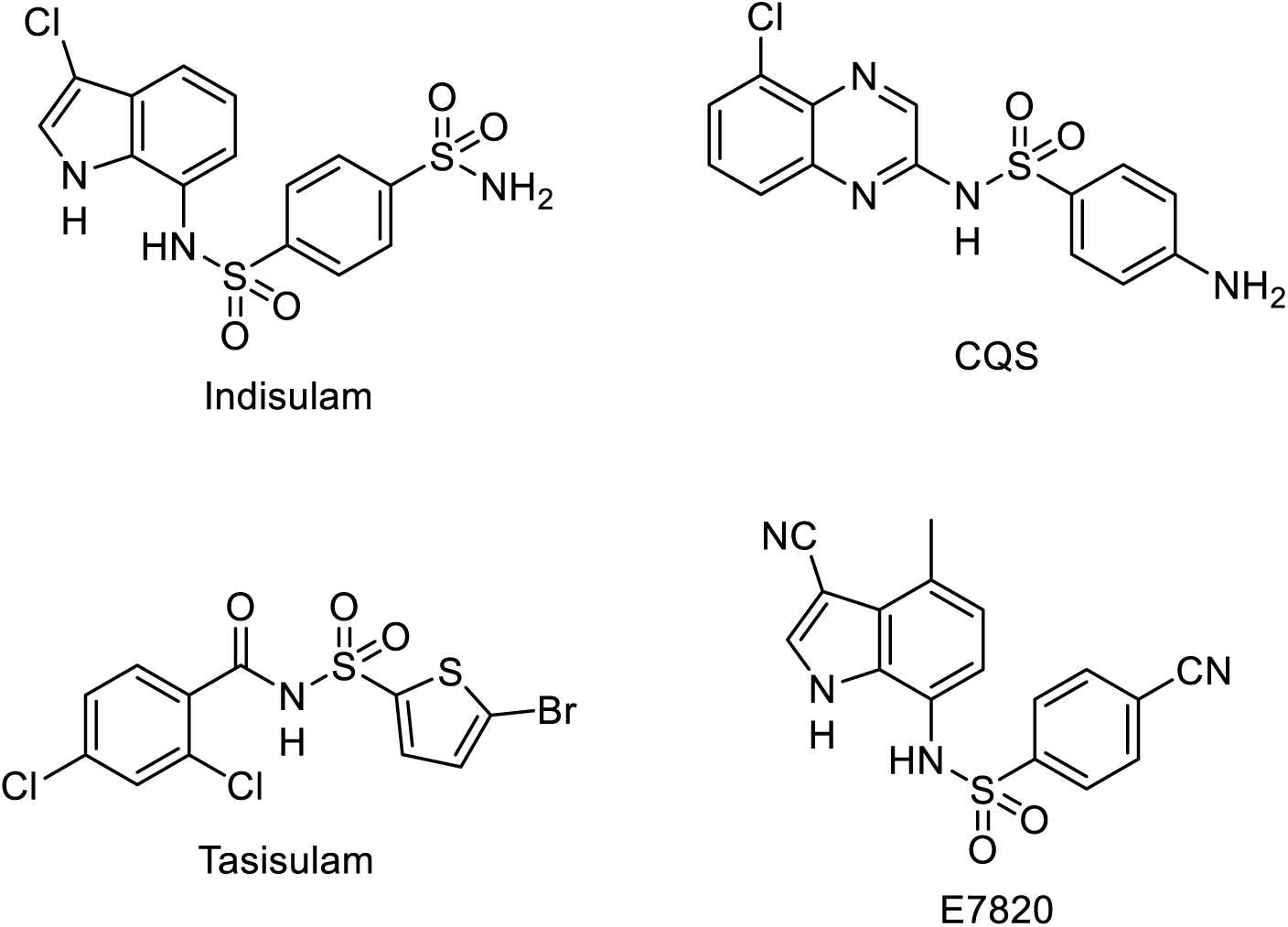
Structures of sulfonamides reported to degrade RBM39 via CRL4^DCAF15^

## Results

### Indisulam as a reprogramable sulfonamide

Our first task was to identify the most modular sulfonamide scaffold that could still retain potency. Additionally an appropriate exit vector from the sulfonamide which did not interupt its binding to DCAF15 would be crucial for creating a bifunctional degrader. Although the sulfonamides have been shown to mediate an interaction between DCAF15 and RBM39, the nature of the interface remains unknown. As such there is no obvious site on the sulfonamides which can be used as a chemical exit vector as in the case of the thalidomides and CRBN. Although the interaction between RBM39 and DCAF15 may not be necessary for reprogramming to target new substrates, in the absence of structural data we chose degradation of RBM39 as a screening method for testing exit vector positions since this phenotype guaranteed that binding to the CRL4^DCAF15^ E3 Ligase was maintained. The photoaffinity probe reported by Uehara *et al.*^18^, ^20^ provided guidance on where indisulam tolerates chemical modifications. For the design of the exit vector of chloroquinoxaline sulphonamide (CQS) we looked at the similarity between its quinoxaline and the indole of indisulam and chose the arylsulfonamide again. Lastly, we modified tasisluam and replaced the thiophene with an aryl group for the synthetic ease and because phenotypic SAR data suggested this was still a potent molecule (reference tasisulam SAR). Hence we synthesized versions of indisulam, CQS and tasisulam with a small linker or a PEG2 linker and examined which still retained their activity (Fig. 2). A *DCAF15^−/−^* cell line was generated in HCT-116 cells^19^ to assess whether the degradation activity of the molecules was indeed DCAF15 dependent. Treating HCT-116 cells with the compunds for 24h showed that the modified indisulam was the least affected by the chemical modifications and was still operating in a DCAF15-dependent manner (Supplementary Fig. 1). Furthermore, the poor performance of the phenyl-modified tasisulam derivative showed that the thiophene is indeed essential for potent RBM39 degradation. Based on these results we went forward with indisulam as the scaffold for testing bifunctional reprogramming.

**Fig. 2:**
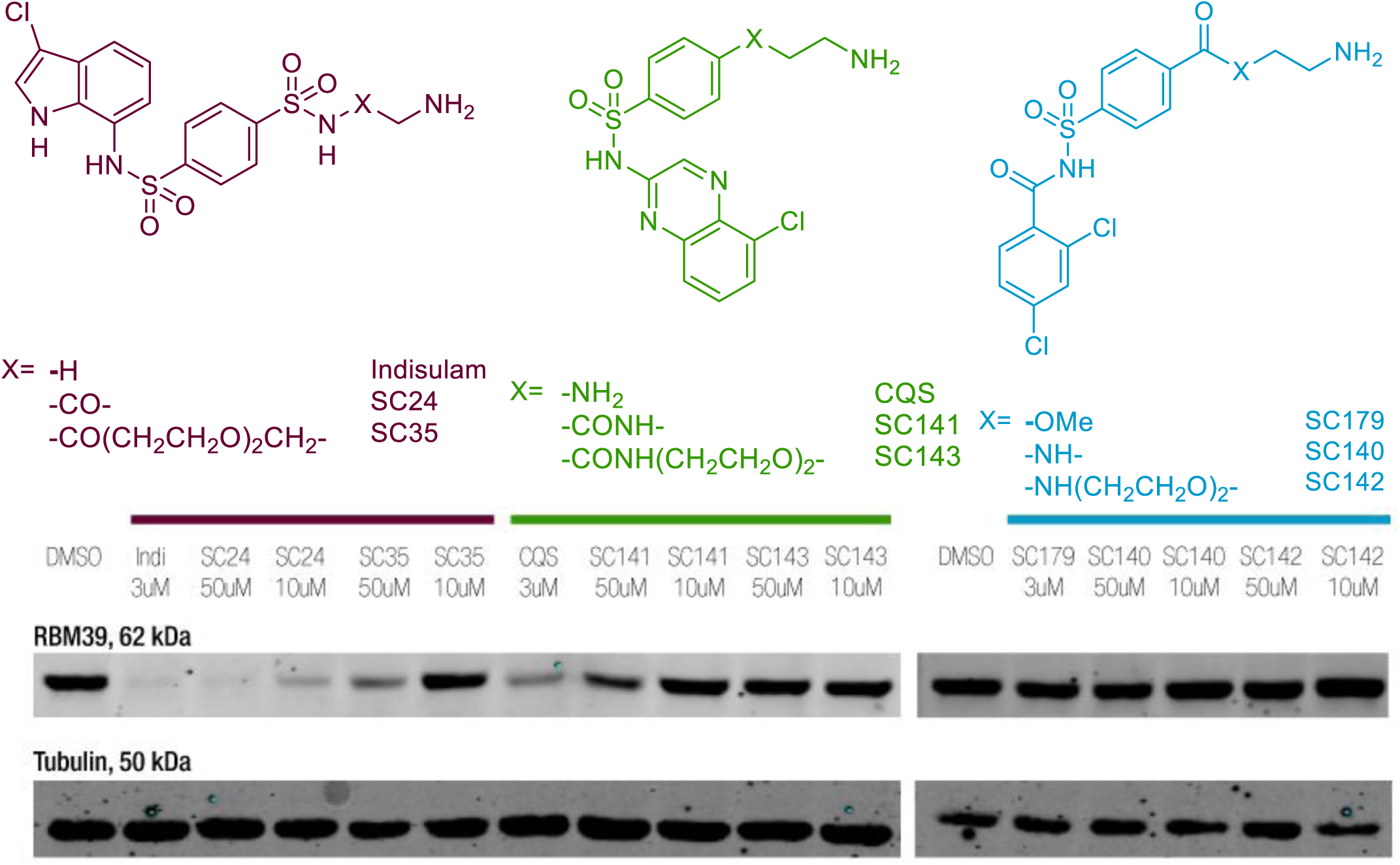
Screening for sulfonamides for the compatibility towards modifications without loss in activity. HCT116 cells were treated with the compounds for 24h at the annotated concentrations

### What is the minimal indisulam scaffold?

A problem with bifunctional PROTACs is that they typically exceed the physicochemical limits for oral bioavailability. Hence it was worthwhile to examine which regions of indisulam were indispensible for activity, so that we could work with the smallest possible pharmacophore. Additionally, the sulfonamide is a relatively inert functional group, and it would be ideal to have a more chemically versatile linking point. Hence we prepared a series of indisulam derivatives that systematically varied the pendant aryl ring or deleted it entirely (Fig. 3). Some of the compounds had been previously tested in a cell viability assay but the mechanism of their toxicity was unkown^21,22^. The results obtained from the treatments show that the arylsulfonamide of indisulam can bear a wide range of groups without losing activity. In contrast, complete deletion of the aryl ring leads to complete abrogation of RBM39 degradation. Treatments in the *DCAF15*^−/−^ confirmed that the activity was indeed DCAF15 dependent. The fact the replacing the sulfonamide of indisulam with an amide (SC169, Fig. 2) did not diminish RBM39 degradation indicated that amide bonds could be tolerated, providing a reliable and versatile chemical handle for building bifunctional molecules.

**Fig. 3:**
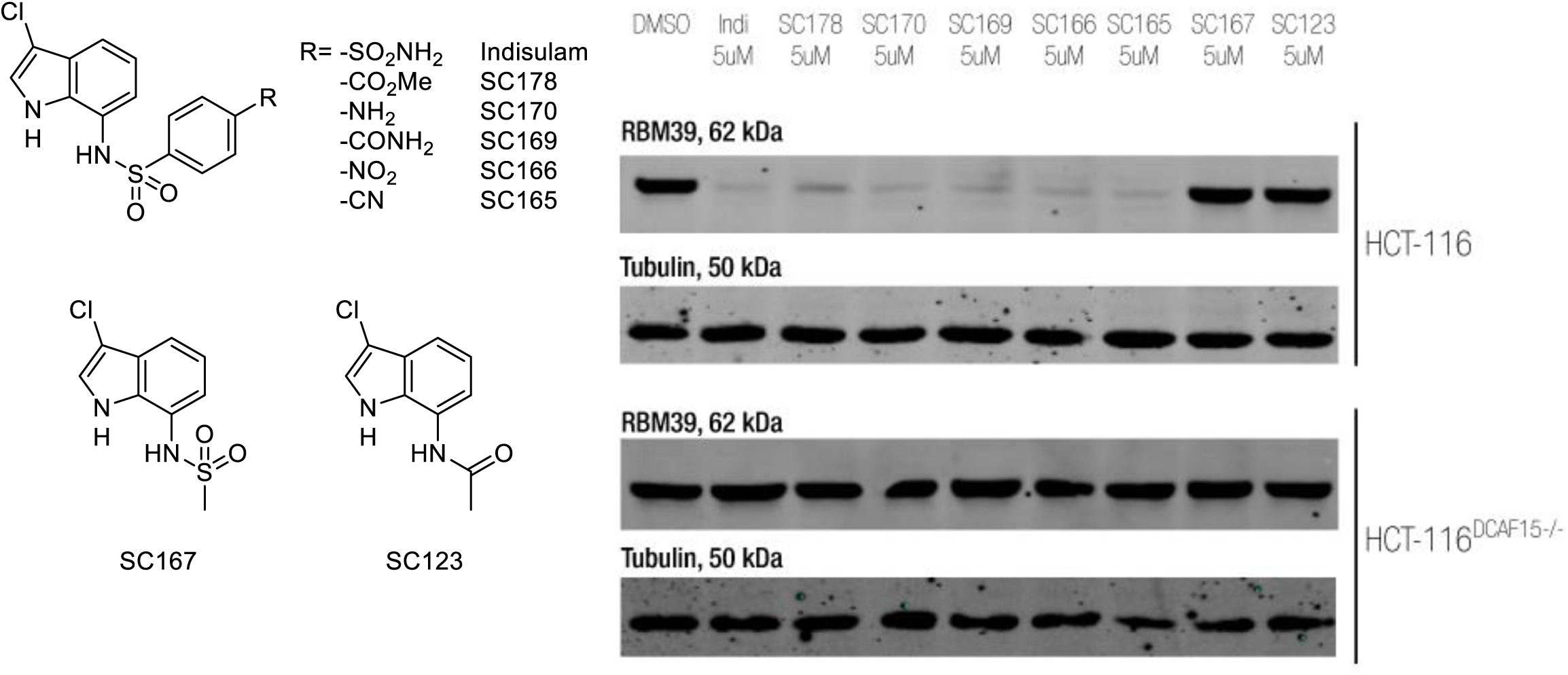
Changing the Indisulam scaffold to see which modifications are tolerated. HCT-116 and HCT-116 DCAF15^−/−^ cells were treated with the compounds for 24h at the annotated concentrations.

### Making bifunctionals against PARP-1 and CRBN

In a next step we made bifunctional molecules by attaching a linker and a small molecule to the indisulam scaffold. A successful bifunctional degrader requires a tertiary interaction between the E3 ligase, the molecule and the protein of interest. Many factors such as the choice of the E3 ligase, the binder of the protein of interest or the linker type and length play a role in this interaction. The linker size and type can be precisely optimized,^23–25^ but medium length PEG linkers (such as PEG3) seem generally to be a good starting point. Since previous work^24,26^ has shown that the success of a given PROTAC is unpredictable, even if the ternary complex is formed, we synthesized two different bifunctional degraders. As a first protein target we chose the Poly (ADP-ribose) polymerase-1 (PARP-1), which has recently been succesfully targeted with the E3 Ligase MDM2.^26^ As a second target we chose lenalidomide, which targets the E3 ligase substrate receptor CRBN. This bifunctional was interesting because it could potentially work in either direction. The CRL4^CRBN^ ligase might degrade RBM39/DCAF15, or the CRL4^DCAF15^ ligase might degrade CRBN.

The molecules were tested in HCT-116 cells for 24h at 50uM, 10uM and 1uM concentrations. Unfortunately with both bifunctionals we saw no degradtion (Fig. 4) at concentrations as high as 50uM. RBM39, however, was still degraded, indicating that the molecules were indeed getting into the cells despite their size and weight.

**Fig. 4:**
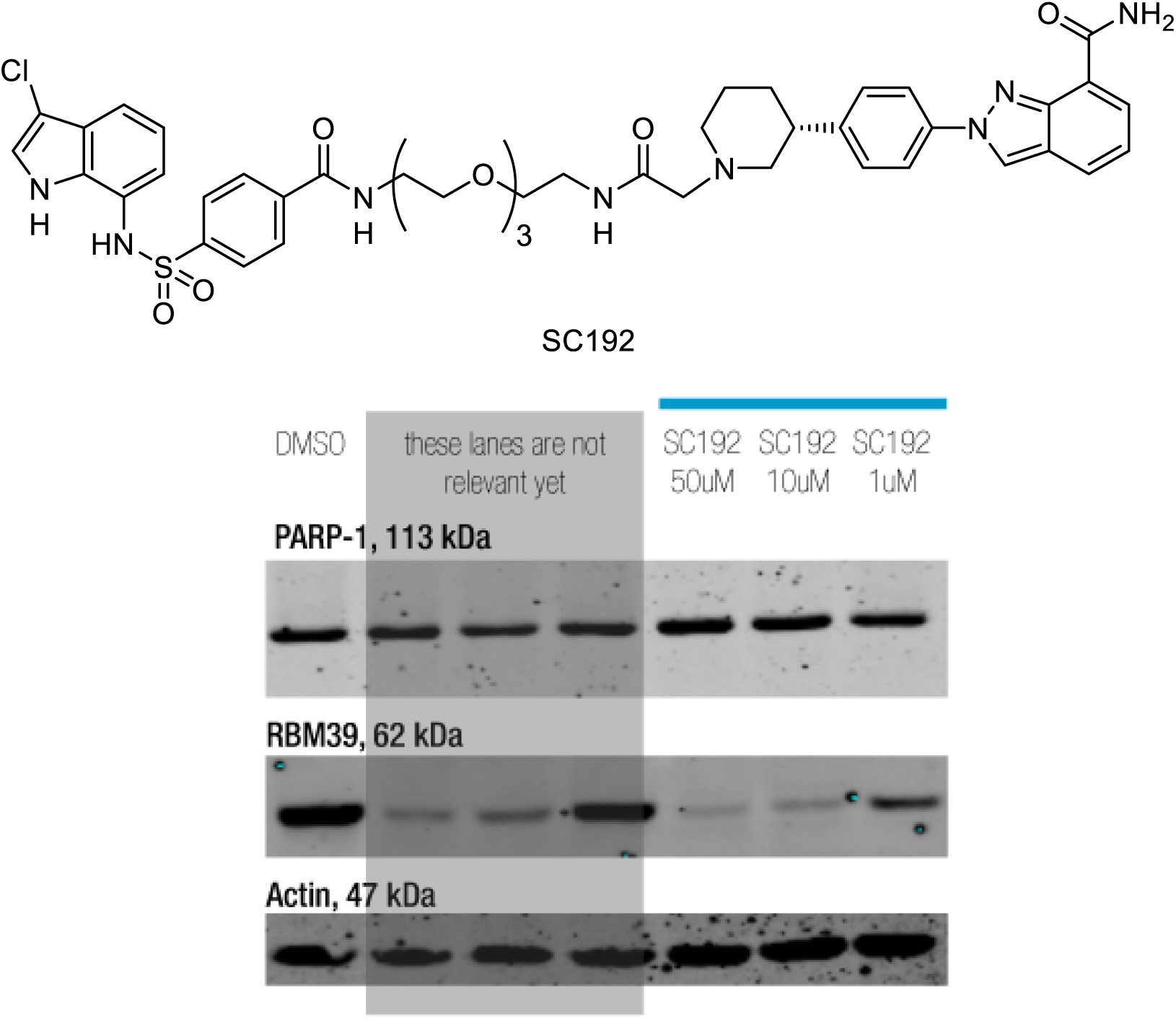
Testing for PARP-1 degradation with the bifunctional molecule SC192. HCT-116 cells were treated with the compounds for 24h at the annotated concentrations

### Binding mode of Indisulam to DCAF15 and RBM39

Although we had assumed a molecular glue-type interaction for RBM39/indisulam/DCAF15, there is in fact no data that illuminates how these three interact. The results reported by Han *et al.*^19^ and Uehara *et al.*^18^ indicate that DCAF15 is the immediate interacting partner of RBM39 in the presence of indisulam *in vivo* and *in vitro* but does not clearly specify as to whether this is direct interaction or an allosteric interaction (Fig. 5). Uehara *et al.*^18^ found that a tritium-labelled variant of indisulam (E7820) is enriched in co-immunoprecipitation experiments only when DCAF15 is present, suggesting weak or rapidly reversible binding to RBM39 in the absence of DCAF15. The lenalidomide derivative (SC193) allows us to investigate this suggestion from a different viewpoint. If indisulam were to have even a weak affinity for RBM39 despite the absence of DCAF15 it might still get degradaed by the CRL4^CRBN^ E3 ligase. Hence we tested SC193 in the *DCAF15*^−/−^ cell lines but saw no degradation of RBM39 at a dosage of 50uM over 24h (Fig. 6). In line with the data provided by Uehara *et al.*^18^ our results imply that an allosteric interaction of RBM39 (Fig. 5c) is unlikely, but we still cannot resolve the other binding options. Although it has been previously shown that CRBN can be degraded via a PROTAC mechanism employing the CRL4^CRBN 27^ or CRL2^VHL 28^ E3 ligases, it could be that this PROTAC is not effective; hence more experiments that speak to the direct binding partner of indisulam would be valuable.

**Fig. 5:**
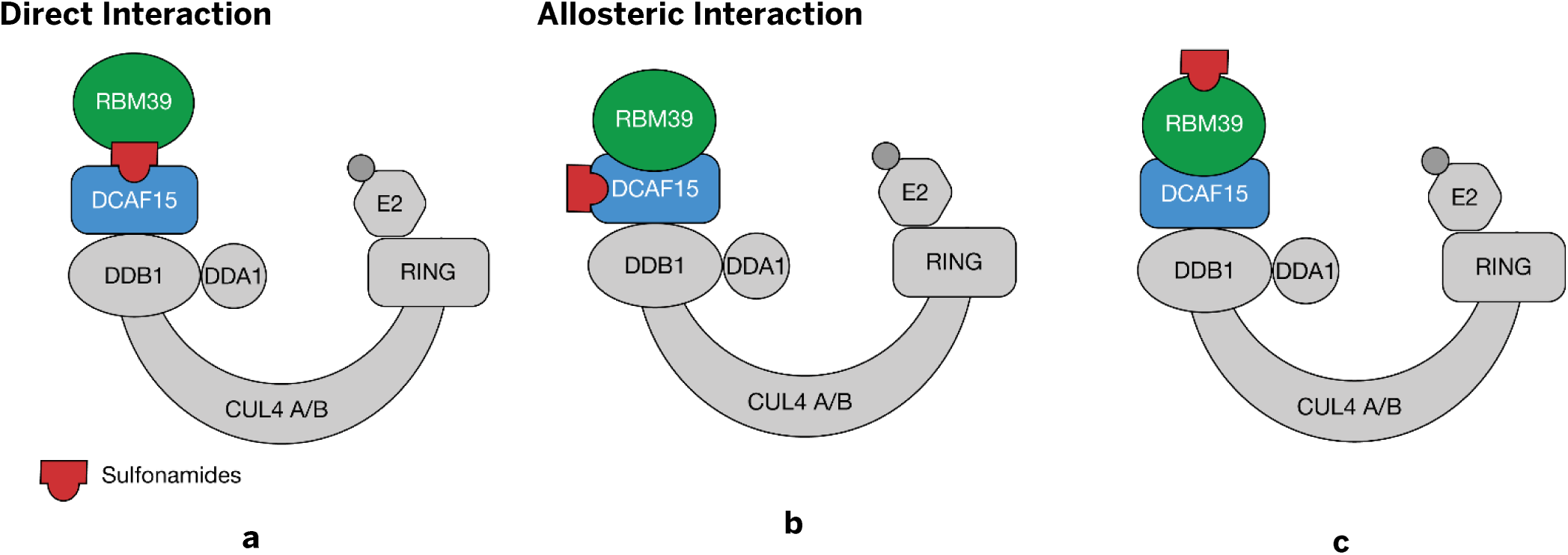
Possible modes of interaction between RBM39 and DCAF15 in the presence of the sulfonamide

**Fig. 6:**
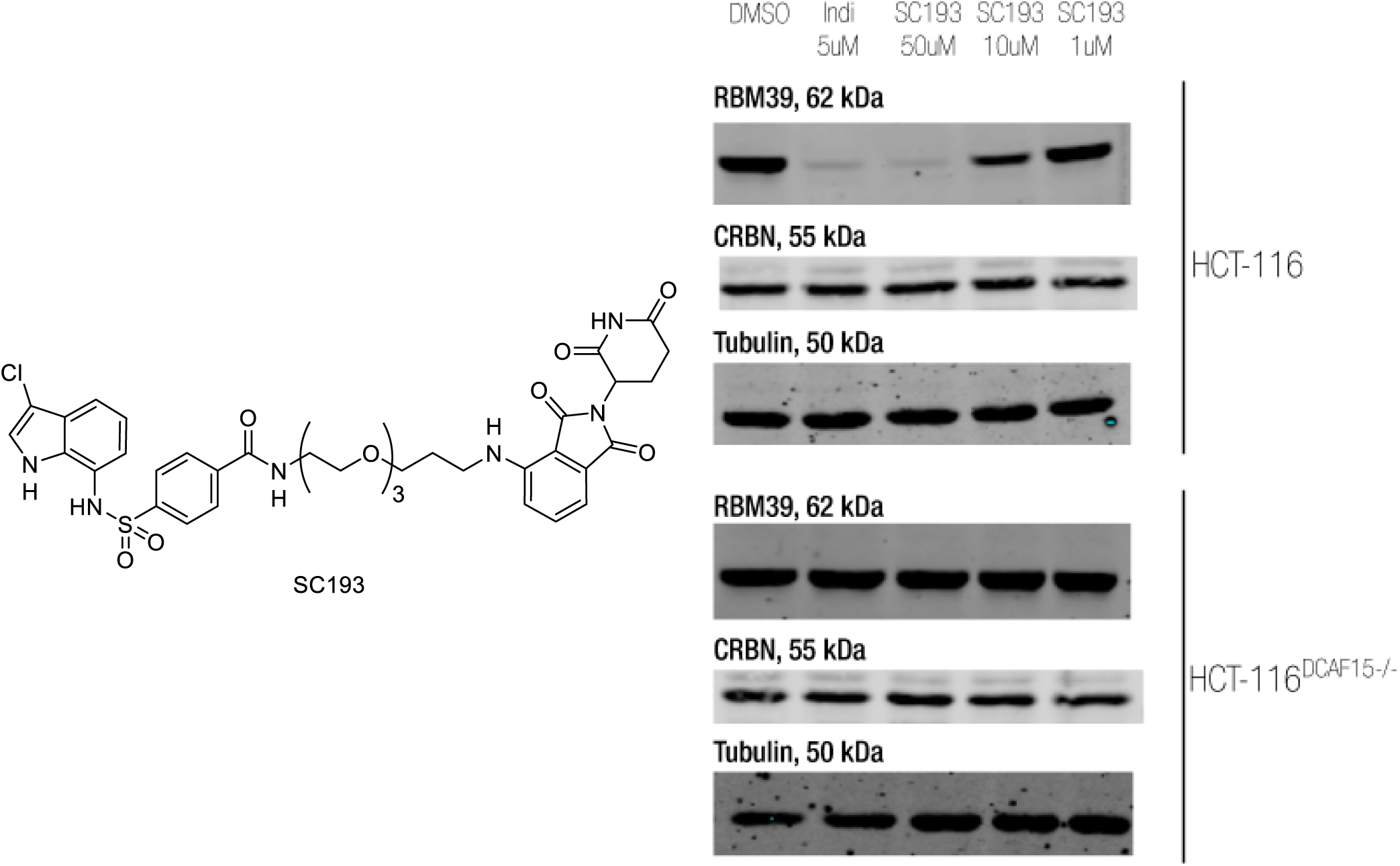
Testing the molecule SC193 in HCT-116 cells for CRBN degradation and in HCT-116 DCAF15^−/−^ cells for RBM39 degradation. Cells were treated with the compounds for 24h at the annotated concentrations

## Conclusion

Our investigations show the indisulam scaffold can tolerate big modifications and still promote CRL4^DCAF15^ dependent degradation of RBM39. This would suggest that there is potential for targeting this E3 ligase for bifunctional degradation, but perhaps different exit vectors need to be analyzed. A clearer picture of the interaction induced between DCAF15 and RBM39 by indisulam would be invaluable for designing bifunctional degraders and we are currently mapping this interaction with chemical proteomics.

---

**Supplementary Fig. 1:**
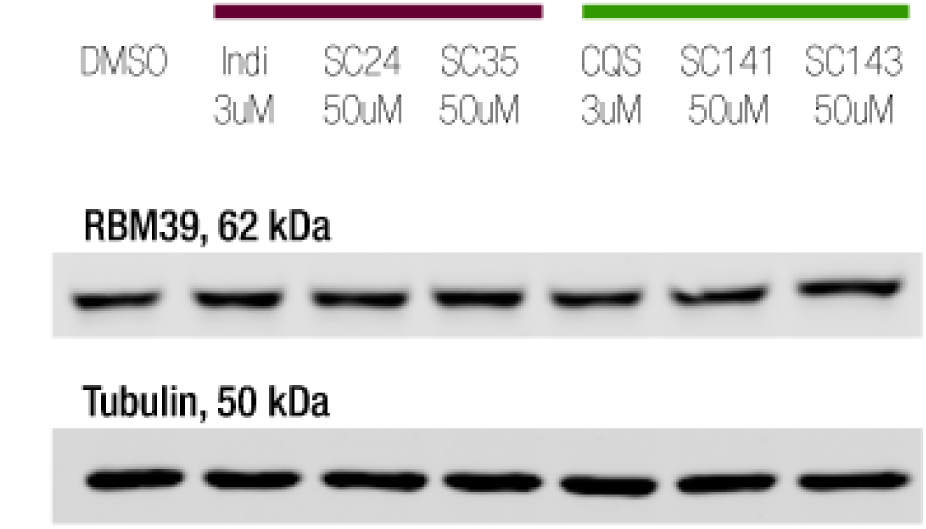
Confirmation of DCAF15 dependent degradation of the modified sulfonamides in Fig. 2. HCT-116 DCAF15^−/−^ cells were treated with the compounds for 24h at the annotated concentrations

